# Age and gender dependence of liver diffusion parameters and the evidence of intravoxel incoherent motion modelling of perfusion component is constrained by diffusion component

**DOI:** 10.1101/2020.08.27.271080

**Authors:** Hua Huang, Cun-Jing Zheng, Li-Fei Wang, Nazmi Che-Nordin, Yì Xiáng J. Wáng

**Author notes:** Corresponding Authors: Dr Yì Xiáng J. Wáng, Department of Imaging and Interventional Radiology, Faculty of Medicine, The Chinese University of Hong Kong, Shatin, New Territories, Hong Kong SAR, China.

## Abstract

**Objectives:** To establish reference values for middle aged subjects and investigate age and gender dependence of liver diffusion MRI parameters.

**Methods:** The IVIM type of liver diffusion scan was based on a single-shot spin-echo type echo-planar sequence using a 1.5-T magnet with 16 *b*-values. DDVD (diffusion-derived vessel density) was the signal difference between *b*=0 and *b*=2 s/mm^2^ images after removing visible vessels. IVIM analysis was performed with full-fitting and segmented-fitting, and with threshold b-value of 60 or 200 s/mm^2^, and fitting started from *b*=2 s/mm^2^. 32 men (age range: 25-71 years) and 26 men (age: 22-69 years) had DDVD and IVIM analysis respectively, while 36 women (age: 20-71 years) had DDVD and IVIM analysis.

**Results:** DDVD had an age-related reduction noted for women. IVIM results of full fitting had good agreement with segmented fitting with threshold *b* of 60 s/mm^2^ results, but less so with results of threshold *b* of 200 s/mm^2^. As age increases, female subjects’ D_slow_ measure had significant reduction, and PF and D_fast_ measure had significant increase. For the age group of 40-55 years, DDVD, Dslow, PF, and Dfast were 12.27±3.90, 1.072±0.067 (10^−3^mm^2^/s), 0.141±0.025, 61.0±14.0 (10^−3^mm^2^/s), and 13.4±3.6, 1.069±0.074 mm^2^/s, 0.119±0.014, 57.1±13.2 mm^2^/s, for men and women, respectively.

**Conclusion:** DDVD measure suggest that aging may be associated with reduction in liver perfusion. Lower D_slow_ measurement can lead to artificial higher PF and D_fast_ measurement, providing the evidence of IVIM modeling of perfusion component is constrained by diffusion component.

## Introduction

Le Bihan *et al.* [1, 2] proposed the principle of IVIM which enables the quantitative parameters that reflect tissue diffusivity (D_slow_,) and tissue microcapillary perfusion (PF, and D_fast_) to be estimated. Recently it was demonstrated that, with careful selection of *b*-values and improved image post-processing, PF and D_slow_ measurement can be quite reproducible, and D_fast_ can be moderately reproducible [3–5]. There are also strong evidences that the relationship between liver DWI signal and *b*-value does not follow bi-exponential decay; instead, it can be better fitted by an addition of very fast component with a tri-exponential decay model [5–12]. However, the fitting of tri-exponential decay model can be quite instable at individual study subject’s level [6, 8, 13]. As an effective alternative to solve this problem, the relationship between liver DWI signal and *b*-value can separated into two parts: part-1 is the signal difference between *b*=0 s/mm^2^ image and the first very low *b*-value image (usually *b*= 1 or 2 s/mm^2^ image), the rest is part-2 and fitted with a bi-exponential decay model [9]. On DWI, blood vessels show a high signal when there is no diffusion gradient (*b* = 0 s/mm^2^) and a low signal even when very low *b* values (e.g., 1 s/mm^2^) are applied. Therefore, for part-1, the signal difference between images when the diffusion gradient is off and images when the diffusion gradient is on reflects the extent of tissue vessel density (referred to diffusion-derived vessel density, DDVD) [5, 14]. For part-2, starting from non-zero very low b-value, the relationship between DWI signal and *b*-value follows a bi-exponential decay [5, 10, 15–18]. It has also been empirically showed that, as compared with the commonly used threshold *b*-value of 200 s/mm^2^ for segmented fitting, the threshold *b*-value of 60 s/mm^2^ to separate slow and fast components performed better in separating normal livers and fibrotic livers [17, 18, 19]. Liver physiological MRI measures may differ in males and females and may differ in young adults and older adults [20–22]. The initial aim of this study was to establish reference IVIM parameter value for middle aged human subjects. We also wanted to know where there is an age and gender dependence of liver DWI parameters among healthy subjects, as this information will be relevant for translating liver IVIM research to clinical practice.

## Material and Methods

The MRI data acquisition was approved by the local institutional ethical committee, and the informed consent was obtained for all subjects. MRI data of healthy participants were acquired during Apr 22, 2019 to Dec 11, 2019 (Table-1). Participants were asked to fast for 6 hours before imaging, and they were scanned twice (scan-1 and scan-2) during the same session whenever possible. The IVIM scan was based on a single-shot spin-echo type echo-planar sequence using a 1.5-T magnet (Philips Healthcare). SPIR technique (Spectral Pre-saturation with Inversion-Recovery) was used for fat suppression. Respiratory-gating was applied in all scan participants. The TR was 1600ms and the TE was 63ms, with one TR per respiratory cycle. Other parameters included slice thickness =7 mm and inter-slice gap 1mm, matrix= 124×97, FOV =375 mm×302 mm, NEX=2, number of slices =6. Images with 16 *b*-values of 0, 2, 4, 7, 10, 15, 20, 30, 46, 60, 72, 100, 150, 200, 400, 600 s/mm^2^ were acquired.

**Table-1.** Study participant demographics*

|  | Males | Females |
| --- | --- | --- |
| DDVD analysis | n=32, age mean:43.3<br>range: 25-71 | n=36, age mean: 43.5<br>range: 20-71 |
| DDVD with two scans** | 31 out of 32 | 36 out of 36 |
| IVIM analysis | n=26, age mean: 42.1<br>range: 22-69 | n=36, age mean: 43.6<br>range: 20-71 |
| ratio with two scans** | 17/26 | 27/36 |

\* 77 healthy volunteers were recruited, and 8 participants' scans (2 males and 6 females) were excluded due to unsatisfactory image quality. Final analysis included 33 males and 36 females, among them 7 males the $b=0$ , 2 images allowed DDVD analysis while their IVIM series suffered from unsatisfactory image quality (thus there were more DDVD results available). One male's $b=0$ images were not acquired, while other images allowed IVIM analysis.
\*\* number of subjects who had repeated two scans and the mean of the two scans were adopted as the results of the subjects.

The measurement of DDVD followed a recent report [14]. ROI for right liver parenchyma was segmented on the *b*=0 s/mm^2^ image (resulting in ROI area of area0) and the *b*=2 s/mm^2^ image (resulting in ROI area of area2) respectively. Since we are interested in liver micro-perfusion, on the *b*=0 s/mm^2^ image we selected a threshold to remove all pixels of vessels which are of bright signal; on the *b*=2 s/mm^2^ image, we selected a threshold to remove all pixels of vessels which are of ‘signal void’. DDVD was calculated according to Eq 1:

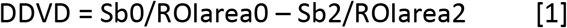

The IVIM analysis for D_slow_ (true diffusion, *D*), PF (perfusion fracture, *f*), and D_fast_ (perfusion related diffusion, *D\**), followed previous descriptions [5, 15–18]. The *b*=2 s/mm^2^ image was used as the starting point for bi-exponential segmented fitting (analysis including *b*=2 s/mm^2^ image is provide in the supplementary document). The signal value at each *b*-value was normalized by attributing a value of 100 at *b*=2 s/mm^2^ (S_norm_=(SI/SI_2_)×100, where S_norm_ is the normalized signal, SI=signal at a given *b*-value, and SI_2_=signal at *b*=2s/mm^2^). Both full fitting and segmented fitting were performed [1, 8, 12], and for segmented fitting the threshold *b*-value to separate the fast component and slow component was 60 or 200 s/mm^2^ [3, 17–19]. For bi-compartmental model, the signal attenuation was modeled according to Eq 2:

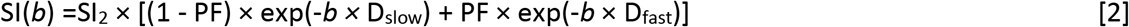

where SI(*b*) and SI_2_ denote the signal-intensity acquired with the *b*-factor value of *b* and *b*=2 s/mm^2^, respectively.

When the two scans for a subject were of good image and fitting quality, the fittings taking the mean of each b-value’s image signal were used; while when one scan was good quality and one scan was unacceptable quality, then the one scan with unacceptable quality was discard (table-1).

## Results

The MRI of thirty-three men (age: 22-71 years) and 36 women’s (20-71 years) were included in final analysis. DDVD measures are presented in Fig-1, and an age-related reduction was noted for women (P=0.01), but not statistically significant for men.

**Fig-1.**
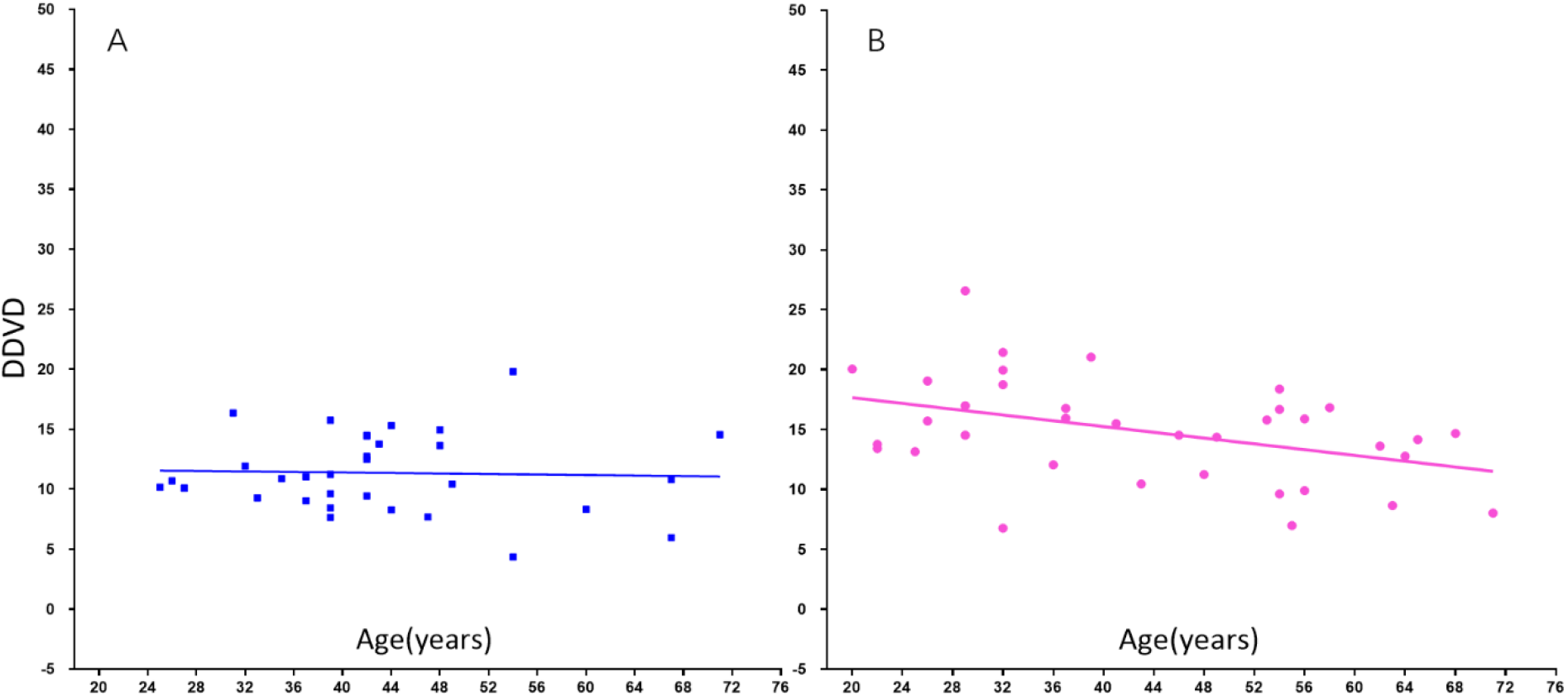

Fig-2 graphically demonstrates IVIM results of full fitting had good agreement with segmented fitting with threshold *b* of 60 s/mm^2^ results, but less so with results of threshold *b* of 200 s/mm^2^. Theoretically, full fitting allows more flexibility and more likely represent physiological measure. Thus, our results further support the adoption of threshold *b* of 60 s/mm^2^ [17–19]. Segmented fitting with threshold b of 60 s/mm^2^ are further presented in Fig-3; and used as primary results for this study.

**Fig-2.**
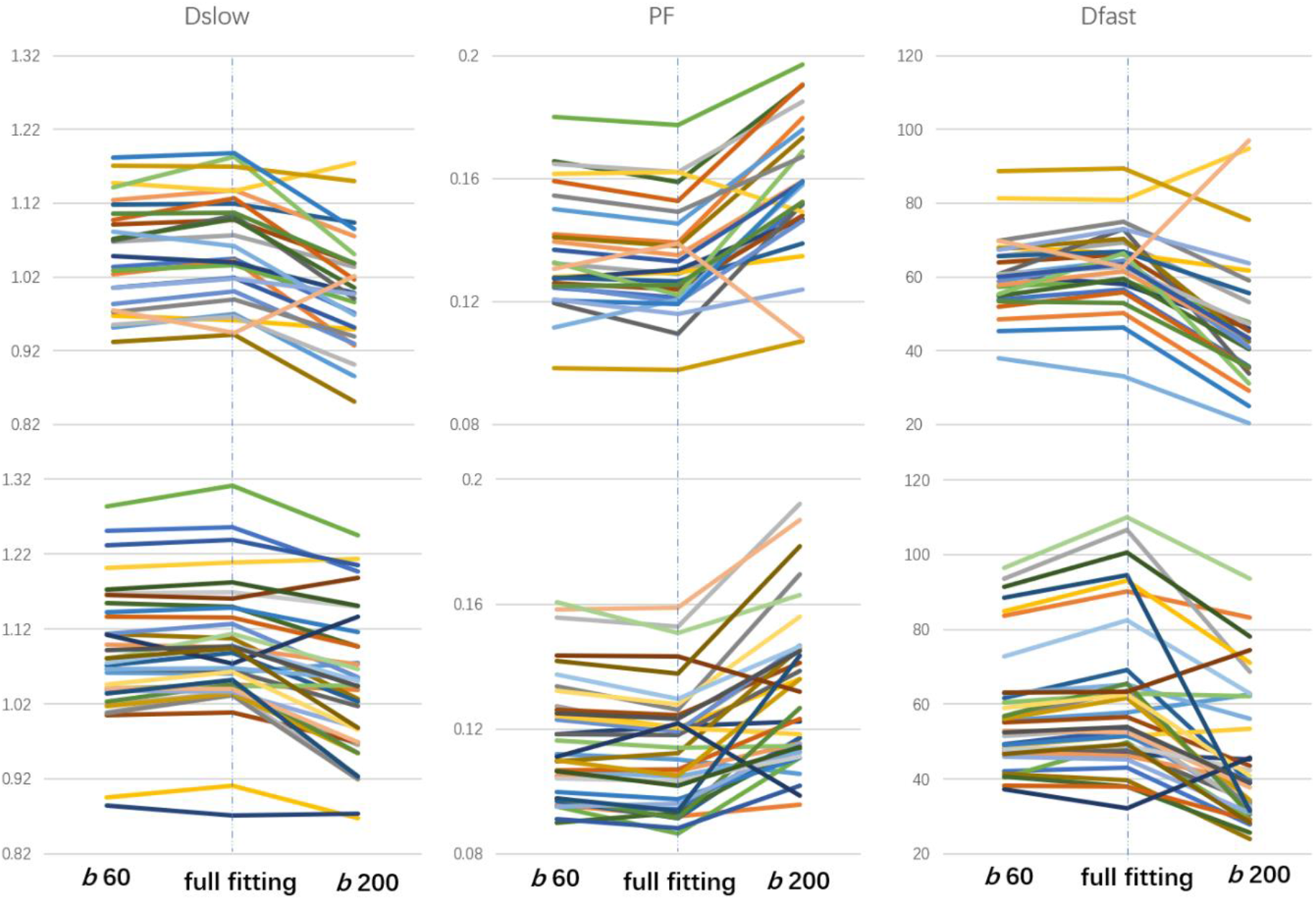
IVIM results of full fitting, segmented fittings with threshhod b=60 s/mm^2^ (*b* 60) and 200 s/mm2 (*b* 200). Upper row for males and lower row for females. Each line represents one subjects.

**Fig-3.**
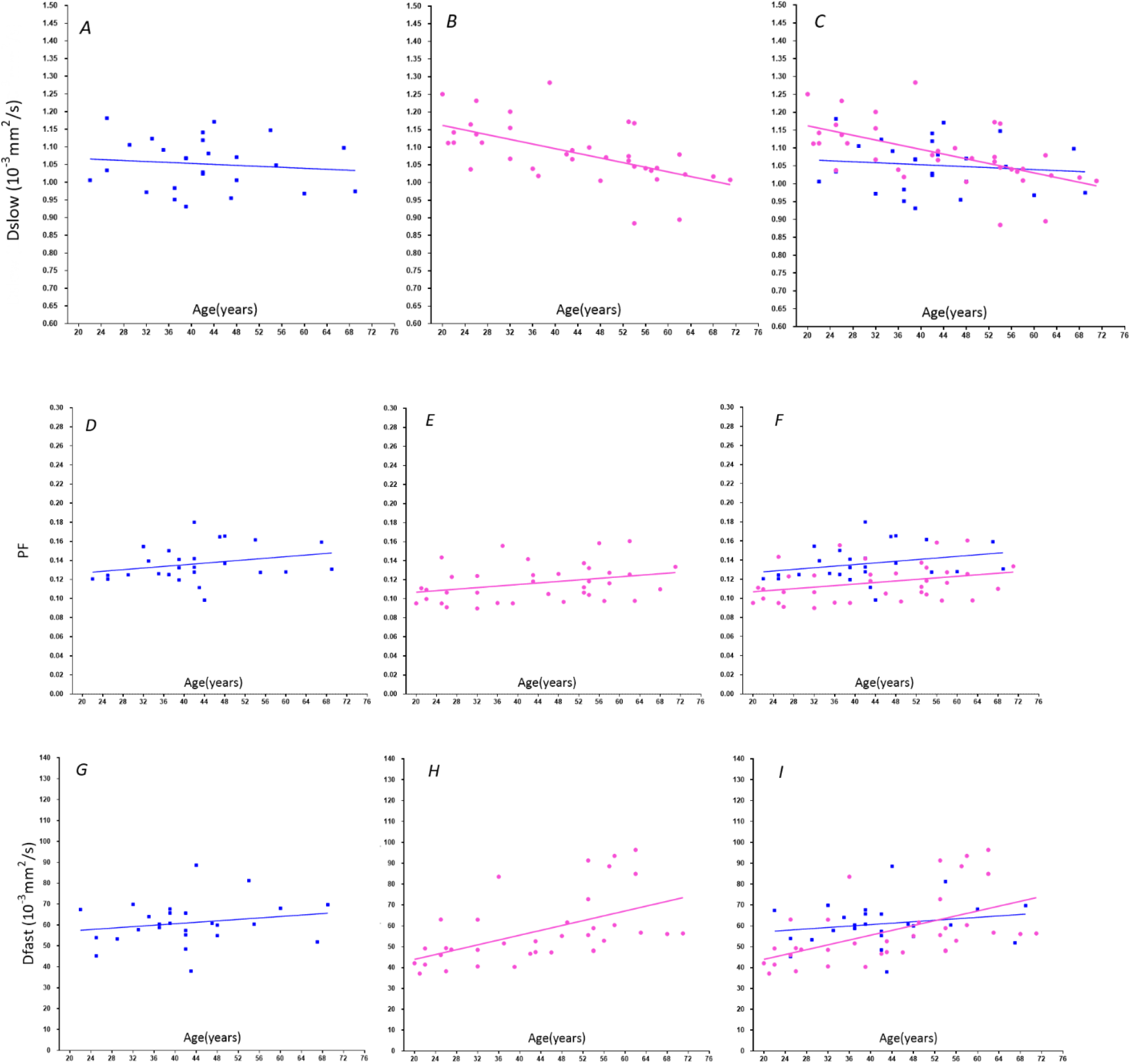
Relationship between IVIM measure and age. Measures were based on segmented fittings with threshhod b=60 s/mm^2^. Blue square: males, red ball: females. C, F, and I show the overlapped males and female’ results.

Fig-3 show, as the age increased, males’ Dslow showed a weak decreasing trend, while female subjects’ Dslow had significant reduction. Males’ PF and D_fast_ showed a weak increasing trend, while women’s PF’s increase trend was close to be significant (P=0.06), and D_fast_’s increase was significant. The linearly fitted lines for males’ and females’ results intercepted at age of 56 yrs for D_slow_ (Fig3C), and at age of 52 yrs for D_fast_ (Fig3I). Compared with young women, young men tended to have lower DDVD and D_slow_ measures and higher PF and D_fast_ measures; while around the age group of 40-55 years, all these measures tended to be similar between men and women. . For the age group of 40-55 years, DDVD, Dslow, PF, and Dfast were 12.27±3.90, 1.072±0.067 (10^−3^mm^2^/s), 0.141±0.025, 61.0±14.0 (10^−3^mm^2^/s), and 13.4±3.6, 1.069±0.074 mm^2^/s, 0.119±0.014, 57.1±13.2 mm^2^/s, for men and women, respectively.

Full fitting results (Fig-4) showed suggest similar findings as described above.

**Fig-4.**
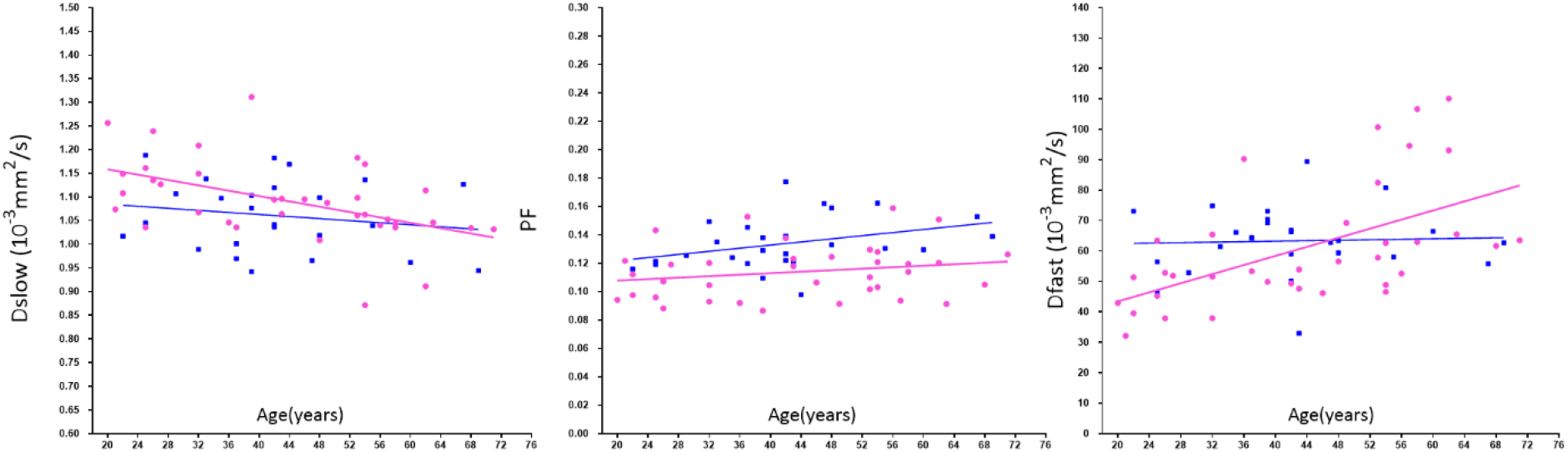
Relationship between IVIM measure and age. Measures were based on full fittings. The trend mirrors results based on segmented fittings with threshhod b=60 s/mm^2^ as shown in Fig-4. Blue square: males, red ball: females.

DDVD and IVIM results for subjects 51 ≤ years of age are shown in Fig-5, showing the above-mentioned trends are also weakly notable before 52 years old (i.e., before women’s menopause age)

**Fig-5:**
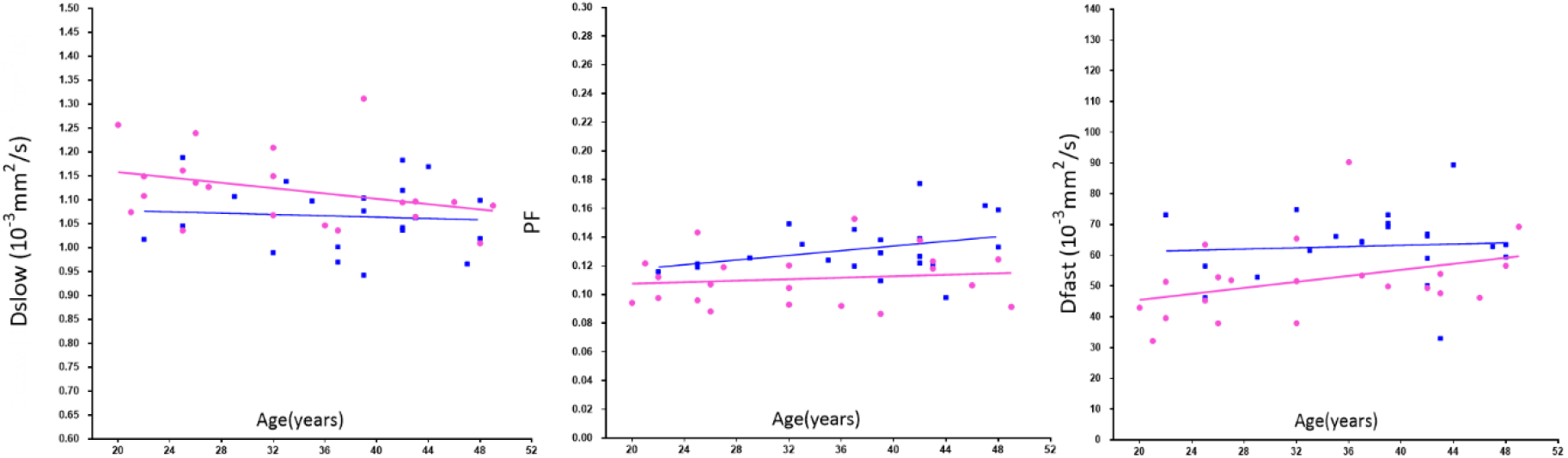
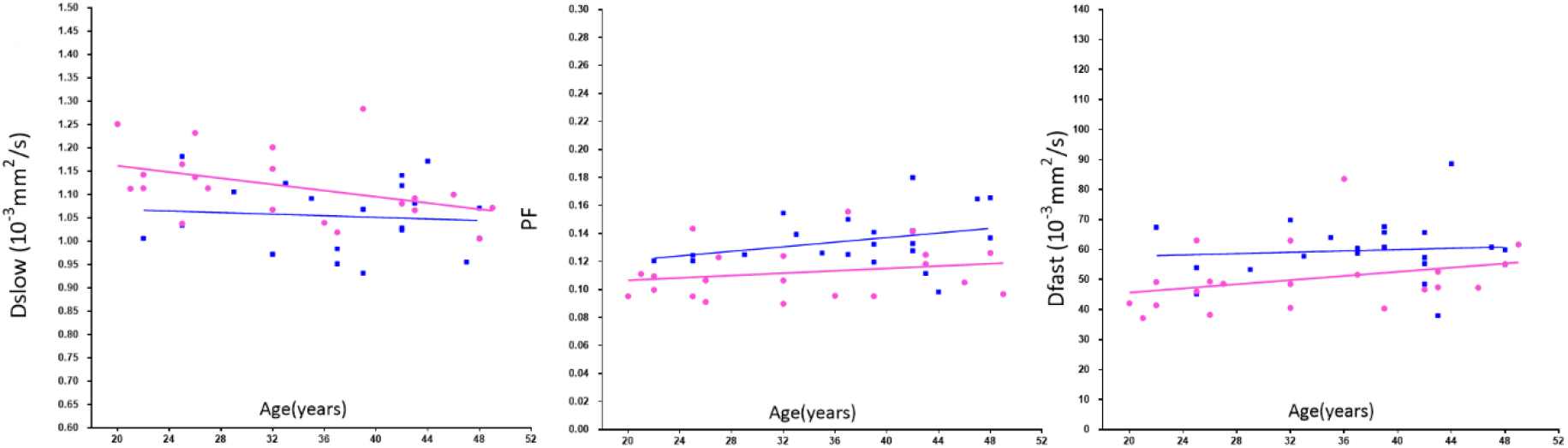
Relationship between IVIM measure and age for subjects less than 52 years. Measures of the upper row are based on full fittings. Measures of the lower row are based on segmented fittings threthold b=60 s/mm^2^. Blue square: males, red ball: females.

## Discussion

The age dependent reduction in liver volume and liver blood flow has been well documented using a variety of technical methods including histology, dye dilution, indicator clearance, and Doppler ultrasound [24–27]. Ours is the first study to demonstrate that the DWI measure shows liver parenchyma had an age-dependent decrease of micro-vessel density (primarily seen in females).

The echo-planar imaging sequence used DWI is subject to susceptibility, and ADC (apparent diffusion coefficient) is known to decrease as a result of the T2* shortening effect [29]. The iron deposition extent in the liver is higher in men than in women [20, 28, 30], and increases over aging, and in women liver iron content increases significantly after menopause [20, 31, 32]. Another possibility, to a lesser degree, would be the influence of liver steatosis. Older age is associated higher prevalent of liver steatosis, and liver steatosis is more prevalent in men than women up to the age of 60 years. Beyond menopause, the prevalence of fatty liver rises sharply in women and exceeds that observed in their male counterparts [33]. Liver steatosis is associated with lower D_slow_ [34]. Intracellular lipid restricts diffusion of water within hepatocytes. Lipid peaks near water are often incompletely suppressed by the chemical shift-dependent suppression techniques used by the DWI sequence; if so, the measured diffusivity may incorporate the diffusion constant of lipid, which is much slower than water [34].

The PF and D_fast_ results gave contradictory results compared with DDVD measure and known physiology of the liver. We hypothesize this was caused by the inability of the IVIM modeling to separate individual parameters, even when full fitting is applied. This study empirically provides the evidence that the quantification of both PF and D_fast_ were constrained by D_slow_, i.e. lower D_slow_ leading to higher PF and D_fast_ measurement, even PF and D_fast_ did not increase or even declined (our IVIM biexponential modeling analysis including *b*=0 s/mm^2^ images demonstrated similar trends, see Fig-6). Considering that D_slow_ is likely affected by tissue iron deposition, and also the measurement for PF and D_fast_ is constrained by D_slow_, D_slow_, PF, and D_fast_ may be called ‘apparent D_slow_’, ‘apparent PF’, and ‘apparent D_fast_’. Chronic liver diseases are often associated with iron overload [35]. In such cases, the real PF and real D_fast_ would be lower than the ‘apparent PF’, and ‘apparent D_fast_’ value. Since chronic liver diseases are usually associated with reduction in perfusion, the DWI measurement sensitivity of perfusion reduction would be attenuated by the liver iron overload. It is also possible that liver iron content affected DDVD quantification, tissues with higher iron content would have higher signal difference between *b*=0 s/mm^2^ images and *b*=2 s/mm^2^ images. Thus, DDVD could be over-estimated in older subjects with higher liver iron content.

**Fig-6:**
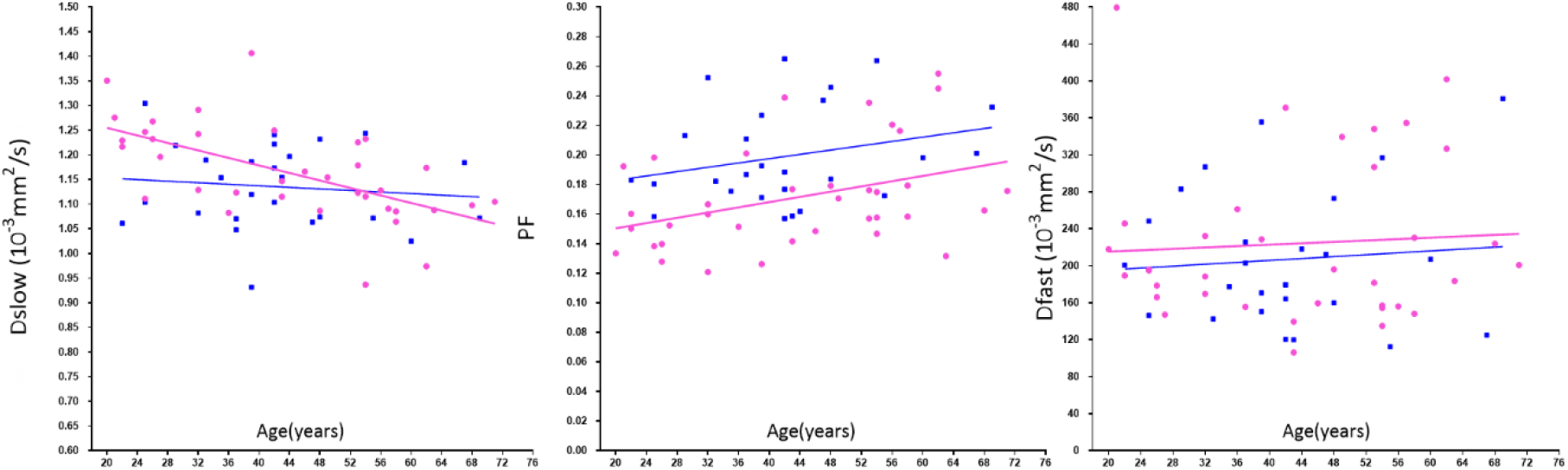
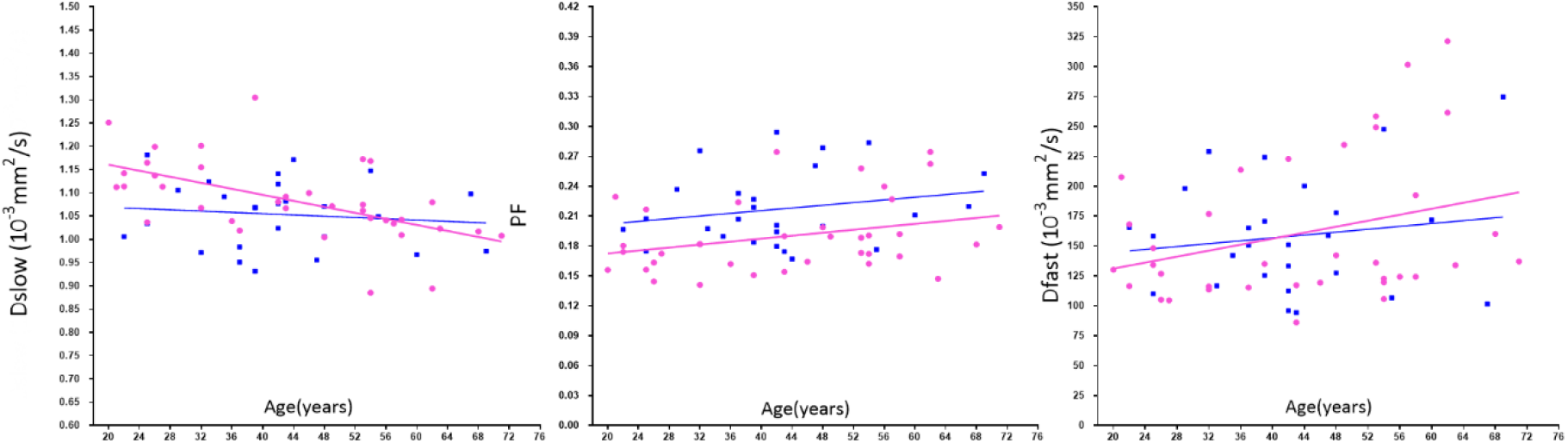
Relationship between IVIM measure and age with IVIM analysis including *b*=0 data. Measures of the upper row are based on full fittings and *b*=0 data included. Measures of the lower row are based on segmented fittings with threthold *b*=60 s/mm^2^ and *b*=0 data included. Blue square: males, red ball: females.

## Abbreviations

DWI: Diffusion-weighted imaging
IVIM: Intravoxel incoherent motion
ADC: apparent diffusion coefficient
DDVD: diffusion-derived vessel density
D_slow_: the intravoxel incoherent motion parameter of tissue ‘true’ diffusion
PF: the intravoxel incoherent motion parameter of perfusion fracture
D_fast_: the intravoxel incoherent motion parameter of perfusion related diffusion. The IVIM analysis for D_slow_ (true diffusion, *D*), PF (perfusion fracture, *f*), and D_fast_ (, *D\**),
ROI: Region-of-interest
Eq.: Equation

## Acknowledgement

This study was partially supported by Hong Kong GRF Project No. 14109218, and the Sanming Project of Medicine in Shenzhen(SZSM201612053).

